# Commonly used insect repellents hide human odors from *Anopheles* mosquitoes

**DOI:** 10.1101/530964

**Authors:** Ali Afify, Joshua F. Betz, Olena Riabinina, Christopher J. Potter

## Abstract

The mode of action for most mosquito repellents is unknown. This is primarily due to the difficulty in monitoring how the mosquito olfactory system responds to repellent odors. Here, we used the Q-system of binary expression to enable activity-dependent Ca^2+^ imaging in olfactory neurons of the African malaria mosquito *Anopheles coluzzii*. This system allows neuronal responses to common insect repellents to be directly visualized in living mosquitoes from all olfactory organs including the antenna. The synthetic repellents DEET and IR3535 did not activate Odorant Receptor Co-Receptor (Orco) expressing olfactory receptor neurons (ORNs) at any concentration, while picaridin weakly activated ORNs only at high concentrations. In contrast, natural repellents (*i.e*. lemongrass oil and eugenol) strongly activated small numbers of ORNs in the mosquito antennae at low concentrations. We determined that DEET, IR3535, and picaridin decrease the response of Orco expressing ORNs when these repellents are physically mixed with activating human-derived odorants. We present evidence that synthetic repellents may primarily exert their olfactory mode of action by decreasing the amount of activating ligand reaching ORNs. These results suggest that synthetic repellents disruptively change the chemical profile of host scent signatures on the skin surface rendering humans invisible to mosquitoes.

## Introduction

Mosquitoes are vectors for many debilitating diseases such as malaria, Zika, dengue fever, and yellow fever. Malaria alone caused an estimated 435000 deaths in 2017^1^. Mosquitoes primarily depend on olfaction, in combination with other senses, to locate their hosts^2,3^. Therefore, targeting the mosquito’s sense of smell using repellent odorants is an effective strategy to prevent them from biting humans. The synthetic compound N,N-diethyl-meta-toluamide (DEET) is the most widely used mosquito repellent in public use since 1957^4,5^. However, DEET has some drawbacks, including high concentrations (∼>30%) are required for it to be effective, an unpleasant odor and oily feeling to some people, and the ability to dissolve some plastics and synthetic rubber^4^. Commercially synthetized alternatives to DEET have been developed (IR3535, picaridin), but these too have similar drawbacks, such as also requiring high concentrations to be effective. In order to improve or identify new repellents, a better understanding of how insect repellents affect a mosquito’s olfactory system is needed. However, the olfactory mode of action of synthetic insect repellents such as DEET, IR3535, and picaridin, as well as natural insect repellents such as lemongrass oil and eugenol, is surprisingly not well understood.

The olfactory system of the *Anopheles gambiae* species of mosquitoes consists of three organs: the antennae, maxillary palps, and labella^2,6,7^. Each of these organs is covered with sensory hairs called sensilla, and each sensillum houses olfactory sensory neurons that may contain one of three types of chemoreceptors: odorant receptors (ORs), gustatory receptors (Grs), and/or ionotropic receptors (IRs). ORs are expressed in the majority of olfactory neurons, and each OR is expressed along with the Odorant Receptor Co-receptor (Orco) to form a receptor complex that is either narrowly or broadly tuned to a variety of host-derived odors^2,6,8^. Grs, specifically Gr22, Gr23, and Gr24, are expressed in sensilla on the mosquito maxillary palps and respond to carbon dioxide^9^. IRs are likely activated by acids and amines^10,11^.

A consensus for how DEET affects the mosquito olfactory system and alters host seeking behavior has not yet emerged. Currently, there are three hypotheses of how DEET affects mosquitoes: 1) DEET directly activates chemoreceptors (ORs, Grs, and/or IRs) on the mosquito antennae, maxillary palps, or the labella to drive repellent behavior (“smell and avoid”)^12–20^; 2) DEET modulates (‘scrambles/confuses’) OR activity in response to odorants^14,15,21–23^; 3) DEET acts directly on the odorant to decrease its volatility and thereby reduces the amount of attractive odorants capable of activating mosquito olfactory receptors (“masking”)^19^. These hypotheses are not necessarily mutually exclusive; DEET may have more than one mode of action.

The mode of action for DEET and other commonly used insect repellents towards *An. gambiae* mosquitoes, which kill more people worldwide than all other mosquito species combined^1^, is the most poorly understood. A lack of understanding is primarily due to the lack of available methods for testing the simultaneous responses of individual olfactory neurons towards DEET or other repellents. Traditionally, insect repellents must be used to individually stimulate each of the ∼750 sensilla using single sensillum recording (a high technical hurdle), or tested against each individual OR ectopically expressed in *Xenopus* oocytes or in the *Drosophila* empty neuron system^20,24^. To address this technical challenge and examine endogenous responses to insect repellents, we generated transgenic *Anopheles coluzzii* (formerly *Anopheles gambiae* M form^25^) mosquitoes in which the calcium indicator GCaMP6f^26^ was expressed in all Orco expressing neurons (genotype: *Orco-QF2*, *QUAS-GCaMP6f*). We used these mosquitoes to re-visit the three leading hypotheses of how DEET and other commonly used insect repellents affect the *An. coluzzii* olfactory system. We found that the natural repellents eugenol and lemongrass oil strongly activate a subset of olfactory receptor neurons, while DEET, IR3535, and picaridin do not directly activate olfactory neurons. These three synthetic repellents instead function as “maskers” of odor-evoked responses. Our data further support the hypothesis that the masking effect of DEET, IR3535, and picaridin in *Anopheles* mosquitoes is not due to direct inactivation of odorant receptors, but instead results from chemical interactions that decrease the amount of activating ligand reaching olfactory receptor targets on the mosquito antennae.

## Results

To examine olfactory responses in all olfactory organs of *An. coluzzii*, we utilized the Q-system of binary expression by generating a mosquito line that contained a *QUAS-GCaMP6f* transgene and crossing this to the validated *Orco-QF2* driver line^7^. The combination of these transgenes directed the expression of the calcium indicator GCaMP6f to all Orco-expressing olfactory neurons. To validate this mosquito model for monitoring odorant-induced olfactory neuron activity, we directly visualized the antennal response to 1 second pulses of six human skin odorants previously shown to activate *An. gambiae* ORs in heterologous expression screens^24^ (Supplemental Fig. 1a). All OR ligands (1-octen-3-ol, 2-acetylthiophene, benzaldehyde, *p*-cresol, 1-hepten-3-ol, and indole) at 1% concentrations elicited olfactory response across the entire antenna (Supplemental Fig. 1). This enabled a rapid method for linking odors to their induced olfactory responses throughout the *An. coluzzii* olfactory system with single-cell resolution. To achieve higher resolution for analysis, we focused on one antennal segment (11^th^ segment) as a representative for antennal neural responses (Fig. 1a; Methods). Fine glass pipette tips were used to flatten down the antenna at basal (segment 1 and 2) and distal segments (12 and 13). Segment 11 was chosen for imaging as it is the most stable distal segment not touched during the preparation. We found that each of the six odorants activated distinct olfactory receptor neurons (ORNs) at the 11^th^ antennal segment (Fig. 1b-d). Together, our results indicated that calcium imaging of olfactory neurons provides a rapid method to interrogate olfactory responses directly in the peripheral olfactory organs of *An. coluzzii* mosquitoes.

**Figure 1.**
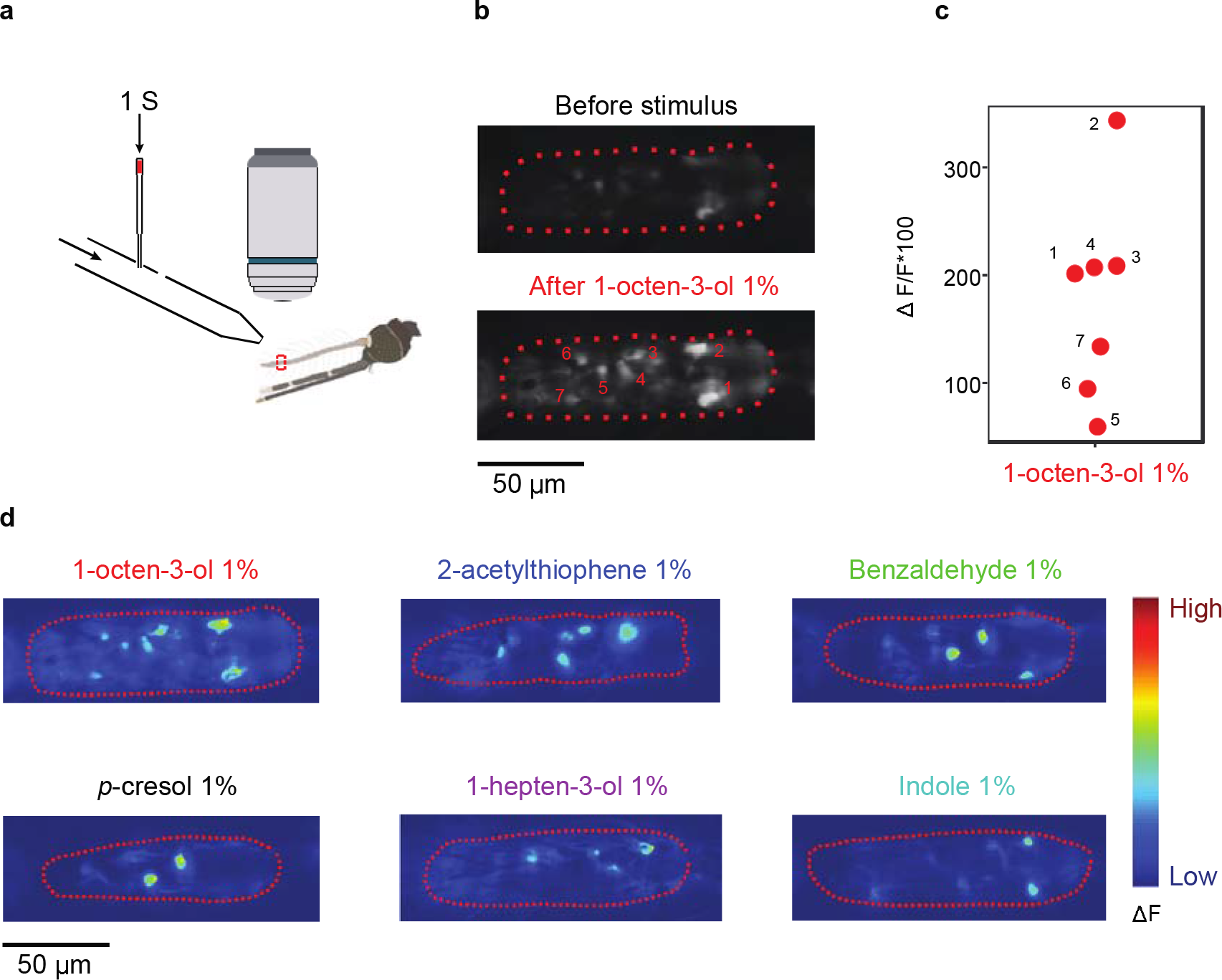
Visualizing odor-dependent activation of *An. coluzzii* antennal olfactory neurons. **a,** Schematic of the calcium imaging setup. A 50x microscope objective images the 11^th^ antennal segment (dashed red rectangle). Arrows indicate the direction of air flow (continuous air, and 1 s air pulse). **b,** Video frames from a calcium imaging recording. Dashed red lines indicate the border of the 11^th^ antennal segment. Numbers identify neurons responding to 1-octen-3-ol at 1%. **c,** ΔF/F*100 values for the neuron responses from the recording in b. **d,** Heatmaps of the responses towards OR ligands at 1%. Dashed red lines indicate the borders of the 11^th^ antennal segment. The heatmap represents arbitrary units.

### Activator and non-activator repellents

A global readout of olfactory neuron function enabled us to investigate how common insect repellents might affect *An. coluzzii* olfactory neurons. We tested two natural repellents (lemongrass oil and eugenol) at 1% concentrations, and three synthetic repellents (DEET, IR3535, and picaridin) at 10% concentrations. We found that the natural repellents lemongrass oil and eugenol elicited strong olfactory responses across the entire antenna, while the three synthetic repellents DEET, IR3535, and picaridin did not elicit any olfactory responses (Supplemental Fig. 2a). We tested all five repellents again with higher resolution imaging at the 11^th^ antennal segment. Lemongrass oil and eugenol at a concentration of 1% strongly activated a subset of ORNs (Fig. 2a) while 10% DEET, IR3535, and picaridin did not activate any ORNs at the 11^th^ segment (Supplemental Fig. 2b).

**Figure 2.**
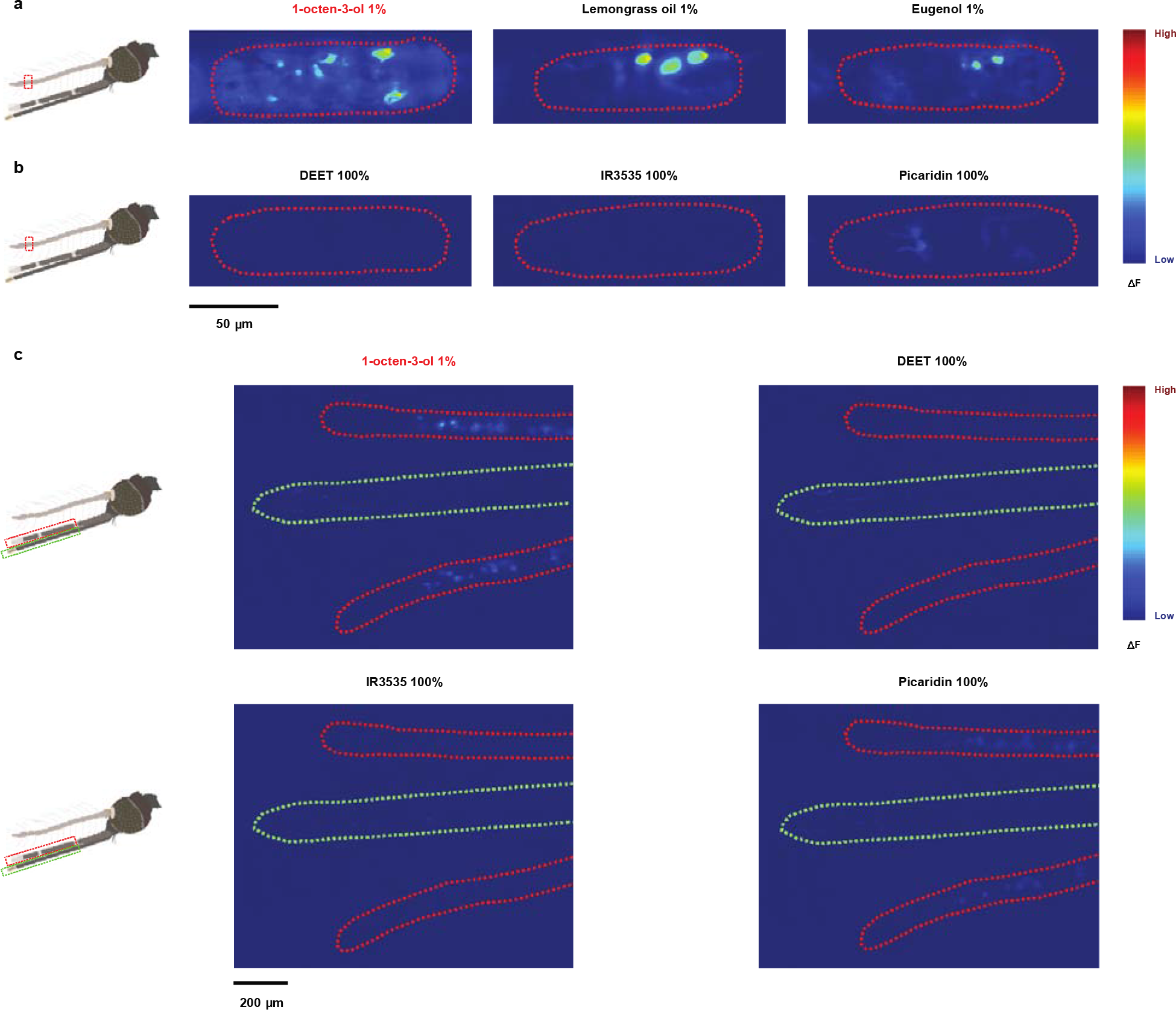
Natural repellents, but not synthetic repellents, strongly activate *Anopheles* olfactory neurons. **a,** Responses at the 11^th^ antennal segment (dashed red line) towards 1% natural repellents lemongrass oil and eugenol. Responses towards 1-octen-3-ol serve as a control stimulus. The heatmap represents arbitrary units. **b,** Responses at the 11^th^ antennal segment (dashed red line) towards 100% synthetic repellents DEET, IR3535, and picaridin. **c,** Responses at the maxillary palps (dashed red line) and proboscis (dashed green line) towards 100% DEET, IR3535, and picaridin, and towards 1% 1-octen-3-ol.

We next asked if higher concentrations of DEET, IR3535, and picaridin would elicit olfactory response in any of the olfactory organs (the antennae, maxillary palps, or labella). There were no olfactory response to DEET or IR3535 at 100% concentrations (Fig. 2b,c). Picaridin at 30% (data not shown) and 100% concentrations elicited a weak response at the antennae, maxillary palps and proboscis. (Fig. 2b,c).

### Synthetic repellents mask odorant receptor ligands

Insect repellents are typically applied directly to human skin and result in a mixture of repellent and human odorants. In this context, DEET might function by altering the olfactory responses to host odorants. Indeed, DEET has been reported to modulate antennal responses towards other odorants in single sensillum recording experiments in *Drosophila*, *Aedes* and *Culex* ^21-23^. In addition, *A. aegypti* olfactory receptors expressed in *Xenopus* oocytes showed an inhibited response towards odorant ligands when mixed with DEET, IR3535, or picaridin^14,15^. We therefore asked if mixing these three repellents individually with known mosquito OR ligands would alter the *An. coluzzii* ORN responses. We found that mixing DEET, IR3535, or picaridin with these activating ligands decreased or “masked” the olfactory neuronal response (Fig. 3, Supplemental Fig. 3a,c,d). In these experiments, each mosquito antenna was tested sequentially with several odorants (OR ligands alone, and mixtures of OR ligands with repellents). These repeated measurements might be correlated within the same animal and were not treated as independent. In addition, there could be an order effect whereby early measurements might affect subsequent measurements. Therefore, we randomized the order of odorants tested, and paired each OR ligands with its respective mixture; *e.g.* OR ligand X was always paired with (precedes or follows) the mixture of OR ligand X + repellent. In addition, we used Linear Mixed Effects regression to account for correlation due to repeated measurements and non-constant residual variation. We found that the masking effect is concentration dependent, where 10% of each repellent showed a significantly stronger masking effect than 1% (Fig. 3b-d, Statistics shown in Supplemental Fig. 3c). Additionally, DEET at 30% masked the response to OR ligands significantly more than 10% (Fig. 3b, Supplemental Fig. 3c). However, there were no differences between the effects of 30% and 10% for both IR3535 and picaridin (Fig. 3c,d, Supplemental Fig. 3c). In addition, there were no differences between the effects of the three repellents when used at the same concentration, except at 10% of DEET and IR3535; DEET showed a significantly weaker masking effect than IR3535 at 10% (Supplemental Fig. 3d). Together, these data indicate that synthetic repellents mask the olfactory responses of OR ligands in a dose-dependent manner.

**Figure 3.**
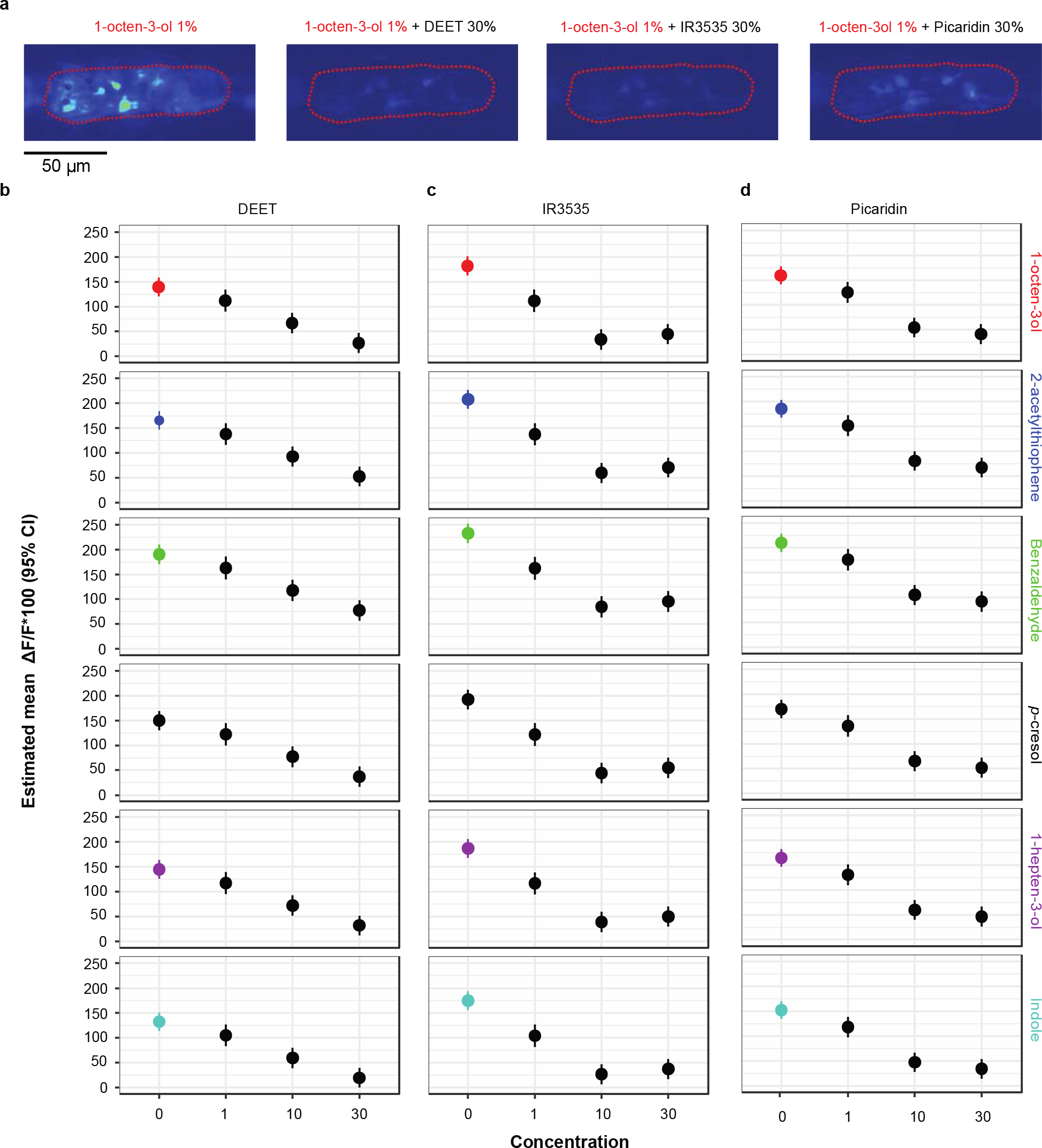
DEET, IR3535, and picaridin mask olfactory responses towards OR ligands. **a,** Heatmaps of the responses towards 1% 1-octen-3-ol and its mixtures with 30% DEET, 30% IR3535, and 30% picaridin. **b-d,** Estimated responses (means and 95% CIs) from Linear Mixed Effect model (LME) towards mixtures of the six OR ligands at 1% with repellents (DEET, IR3535, and picaridin) at 0% (OR ligand alone), 1%, 10%, and 30% (n=15-17 animals for each condition of 0% repellent, n=5-7 animals for all other conditions, 1-7 responding olfactory neurons/animal). All raw data are reported in Supplemental Fig. 3a.

We also asked whether a potentially more potent repellent could be produced by mixing an activator repellent with a masker repellent. We found the ability of activator repellents to stimulate olfactory neurons could also be suppressed by masker repellents; mixing eugenol with DEET, IR3535, or picaridin strongly decreased the eugenol-alone olfactory response. However, the response to lemongrass oil was only partially decreased (Supplemental Fig. 3b). This suggests that a neuron-activity profile approach might be useful in potentially identifying effective repellent combinations.

### Olfactory Masking Requires Chemical Interactions

We sought to understand the mechanism by which repellent masking might occur in *An. coluzzii*. We hypothesized it might occur by one of two potentially overlapping mechanisms. First, olfactory masking could occur at the odorant receptor level, whereby the repellent binds to an odorant receptor complex and prevents its activation by other odorants^14,15,21–23^. Second, olfactory masking might occur at the chemical level by which the repellent reduces the volatility of an odor, resulting in decreased neuronal responses^19^. To determine whether masking occurs at the odorant receptor level, we modified how the repellents and OR ligands were delivered to the mosquito antenna in our system. Instead of delivering a 1 second pulse of either the OR ligands or the repellent and OR ligands mixture, we first delivered a 3 second pulse of the repellent. This allowed the repellent to arrive at the antenna before the OR ligands, and potentially inhibit olfactory receptor complexes. During the last second of repellent odor delivery, we separately delivered a pulse of 1-octen-3-ol into the repellent odor stream (Fig. 4a). If masking occurs at the odorant receptor level, we predicted the repellent would bind to the odorant receptor and inhibit its response towards the delayed OR ligand stimulus. This was not observed. Instead, we found no difference between the olfactory response to 1-octen-3-ol when delivered after a pre-stimulation with each of the three masker repellents and the response when delivered after the control odor paraffin oil (Fig. 4a, Supplemental Fig. 4 a,b). All olfactory responses remained higher than the response to the 1-octen-3-ol mixed with the repellent (Fig. 4a, Supplemental Fig. 4 a,b). This suggests that olfactory masking in *An. coluzzii* does not occur at the receptor level, but more likely at a chemical level.

**Figure 4.**
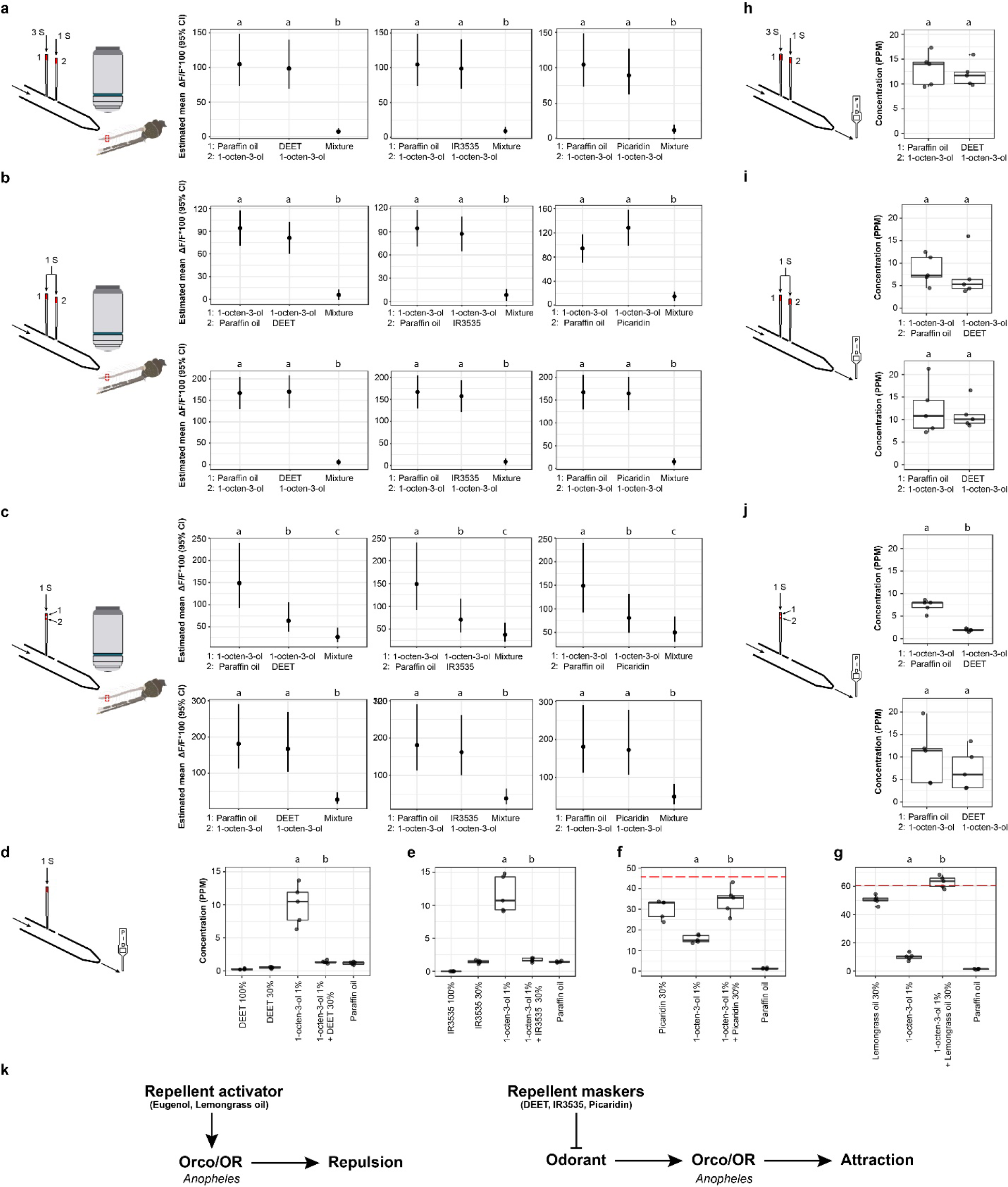
Repellent maskers in *Anopheles* decrease olfactory responses by reducing the volatility of OR ligands, and not by inhibiting odorant receptors. **a,** Estimated responses (means and 95% CIs) from LME towards a 1 s pulse of 1% 1-octen-3-ol occurring during the last second of a 3 s pulse of paraffin oil, 30% DEET, 30% IR3535, or 30% picaridin, compared to the response towards physical mixtures of 1% 1-octen-3-ol with 30% DEET, 30% IR3535, or 30% picaridin. The numbers next to odorant names indicate the position of the odorants in the Pasteur pipette(s) as shown in the schematic. **b,** Estimated responses (means and 95% CIs) from LME towards a 1 s pulse of 1% of 1-octen-3-ol in the first position or the second position simultaneously delivered with a 1 s pulse of paraffin oil, 30% DEET, 30% IR3535, or 30% picaridin, compared to the response towards physical mixtures of 1% 1-octen-3-ol with 30% DEET, 30% IR3535, or 30% picaridin. **c,** Estimated responses (means and 95% CIs) from LME towards a 1 s pulse of 1% 1-octen-3-ol when applied on the upper filter paper or the lower filter paper with paraffin oil, 30% DEET, 30% IR3535, or 30% picaridin in the same Pasteur pipette, compared to the response towards physical mixtures of 1% 1-octen-3-ol with 30% DEET, 30% IR3535, or 30% picaridin. For a-c, n=5 animals for each condition (1-6 responding neurons/animal), conditions denoted with the same letter were not significantly different (P > 0.05, LME model with Wald approximation) Pairwise comparisons between subsequent concentrations are shown in Supplemental Fig. 4b,d,f. Corresponding raw data for a-c are reported in Supplemental Fig. 4a,c,e. **d-g,** Total concentrations (tested by the PID) of odorants released from Pasteur pipettes containing single odorants or their mixtures. Box plots represent the median and 25^th^-75^th^ percentiles. Dashed red line in f indicates the calculated sum of the mean concentrations released from the 1-octen-3-ol and picaridin pipettes. Dashed red line in g indicates the calculated sum of the mean concentrations released from the 1-octen-3-ol and lemongrass oil pipettes. The 10.6 eV PID did not detect DEET or IR3535. **h,** Total concentrations released from the 1% 1-octen-3-ol pipette following a 3 s pulse of 30% DEET or paraffin oil. **i,** Total concentrations released from the 1% 1-octen-3-ol pipette in the first position or the second position when a 1 s pulse of 30% DEET or paraffin oil were used simultaneously. **j,** Total concentrations released from 1% 1-octen-3-ol applied on the upper filter paper or the lower filter paper, while 30% DEET or paraffin oil are applied in the same pipette. For d-j, n=5 experiments. Concentrations denoted with different letters were significantly different (Welsh Two Sample t-test, P < 0.05). **k,** Model for the effects of insect repellents on olfactory responses in *Anopheles* mosquitoes. Natural repellents (Eugenol and lemongrass oil) activate a subset of ORs leading to repulsion of *Anopheles* mosquitoes. Synthetic repellents (DEET, IR3535, and picaridin) interact with OR ligand odorants to mask the attraction of *Anopheles* mosquitoes.

We next asked if repellent masking occurs only to odorants mixed with repellents in the liquid phase (as when on human skin) or might also occur during mixing as volatiles. To answer this question, we delivered the two odorants separately and simultaneously through a Y-tube to allow their molecules to mix in the headspace inside a long pipette directed at the antenna (Fig. 4b). In this setup, there was no difference between the response to 1-octen-3-ol when delivered separately from the repellent and when 1-octen-3-ol was delivered with the control odor paraffin oil; the position of the stimulus pipette relative to the repellent pipette likewise had no effect on altering odorant responses (Fig. 4b, Supplemental Fig. 4c,d). The olfactory responses were significantly higher than the response to 1-octen-3-ol when it was physically mixed with a repellent (Fig. 4b, Supplemental Fig. 4c,d). To confirm that physical mixing is required for masking, we applied 1-octen-3-ol and a repellent on two separate filter papers inside the same Pasteur pipette (Fig. 4c). In this setup, the odorants from the upper filter paper would pass by the lower filter paper as they travel towards the antennae. We found no repellent masking effect when the repellent was on the upper filter paper, but the response to 1-octen-3-ol was significantly reduced when DEET, IR3535, or picaridin were applied to the lower filter paper (Fig. 4c, Supplemental Fig. 4e,f). This second setup mimics situations in which a masker repellent is applied to clothing, which may allow the activating OR ligand to mix with the repellent on their way towards the mosquito antenna. Nonetheless, the olfactory response in the non-mixed condition remained significantly higher than the response to 1-octen-3-ol when it was physically mixed with DEET, IR3535, or picaridin (Fig. 4c, Supplemental Fig. 4e,f). Altogether, these data suggest that masking occurs most effectively when the OR ligand and synthetic repellent are physically mixed, but can also occur to lesser degrees when such ligands travel over a repellent solution that might trap these molecules.

### Masker repellents reduce the concentrations of OR ligands reaching the antenna

The calcium imaging experiments indicate that masker repellents reduce neuronal responses to the panel of OR ligands we have tested. We hypothesized this neuronal effect occurs due to a reduction in the volatility of the odorants we tested which results in fewer ligand molecules reaching the antennae capable of activating olfactory neurons^19^. To test this hypothesis, we used a photoionization detector (PID) to measure the concentrations of odorants that reached the antenna during the different imaging experiments (Fig. 4d-j). The PID measures the total concentration of odorant molecules in air but does not identify these odorants. We found that DEET and IR3535 were likely not detectable by the 10.6 eV PID (Fig. 4d, e). The mixtures of 1-octen-3-ol with 30% DEET or 30% IR3535 showed significantly lower concentrations of odorant molecules than 1-octen-3-ol alone (Fig. 4d, e). This supported the hypothesis that physically mixing the OR ligand with DEET or IR3535 resulted in a lower concentration of that test odorant reaching the antenna. On the other hand, picaridin was strongly detected by the PID, and when 1-octen-3-ol was mixed with picaridin, the mixture showed a concentration that was significantly higher than 1-octen-3-ol alone (Fig. 4f), but not significantly different than picaridin alone (Fig. 4f). Nonetheless, the concentration detected from the mixture was lower than the expected sum of the mean concentrations of the two individual odorants (Fig. 4f), suggesting that picaridin was likely decreasing the levels of volatile 1-octen-3-ol reaching the PID. As a control, we tested 1-octen-3-ol mixed with an activator repellent (lemongrass oil), and found the mixture showed odorant concentrations equal to the expected sum of the individual components (Fig. 4g).

Finally, we used the PID to determine if decreased volatility might also underlie the results obtained under the three modified odorant delivery methods (Fig. 4h-j). We found the concentration of 1-octen-3-ol was unchanged when delivered after a pre-stimulation with DEET or paraffin oil (Fig. 4h). The concentration of 1-octen-3-ol similarly did not change when delivered simultaneously (but not-mixed) with DEET (Fig. 4i). The concentration of 1-octen-3-ol significantly decreased when applied on the upper filter paper in the same Pasteur pipette with DEET on the lower filter paper (Fig. 4j). These PID experiments support our hypothesis that the masking effect observed during calcium imaging experiments was due to a lower concentration of the OR ligand we screened reaching the antenna when the OR ligand was physically mixed with or trapped by a masker repellent. The differential effects of the three masker repellents on olfactory responses likely reflects their chemical differences in altering OR ligand volatilities.

Our calcium imaging experiments support two modes of action for olfactory repellents in *An. coluzzii* (Fig. 4k): 1) Natural repellents such as eugenol and lemongrass oil activate subsets of Orco/OR-expressing olfactory neurons to guide mosquito repulsion, and 2) synthetic repellents do not activate Orco/ORs directly, but instead chemically interact with OR ligands to prevent them from reaching the mosquito antenna. Chemical masking by synthetic repellents may therefore act directly on the skin surface to dramatically alter the chemical profile of human volatiles released into the environment, potently disrupting mosquito olfactory attraction.

## Discussion

By globally monitoring olfactory neuron responses to odors, we present evidence that adult *An. coluzzii* Orco-expressing olfactory neurons do not directly respond to three of the most commonly used synthetic repellents (DEET, IR3535, and picaridin). These findings differ from studies exploring DEET perception in *Culex* and *Aedes* mosquito species. *Culex quinquefasciatus* mosquitoes encode an odorant receptor (*CqOR136*) activated by DEET, IR3535 and picaridin when expressed with *CqOrco* in *Xenopus* oocytes^19,20^. Although a DEET receptor remains to be identified in *Aedes aegypti* mosquitoes, *orco* mutant behavioral studies suggest that Orco-expressing olfactory neurons are likely necessary for DEET-based responses in the presence of human odor^16^. Interestingly, *An. coluzzii* larvae behaviorally respond to DEET in water^27^; however, DEET detection in this context might be mediated by a larval-specific OR or via non-olfactory neurons.

Calcium imaging is a powerful approach to simultaneously visualize the odor-induced activity of many olfactory neurons, but it does have technical limitations. For example, calcium imaging studies may not be able to detect olfactory neurons only weakly activated by DEET or other repellents; however, in the current study, even 100% DEET (a concentration 3-fold higher than commonly effective) failed to activate olfactory neurons, suggesting that any neurons missed by our study would likely express only low affinity DEET-receptors. Calcium imaging may also poorly detect neuronal inhibition (potentially visualized as a decrease in basal GCaMP6f fluorescence); nonetheless, the effects of neuronal inhibition on odor-induced activities would have been easily detectable (Fig. 4), and their absence suggests any direct inhibitory effect is negligible. Finally, in the current work, GCaMP6f is expressed specifically in Orco-expressing neurons, and will not label olfactory neurons that express ionotropic or gustatory receptors. Even though only Orco expressing neurons in other mosquitoes have been reported to respond to DEET, it is possible that DEET, IR3535, and picaridin might be sensed by *An. coluzzii* through non-Orco olfactory pathways.

DEET, IR3535 and picaridin likely exhibit multiple overlapping modes of action in preventing mosquito bites. Their ability to function as chemical maskers undoubtedly translates into their function in masking attraction of humans to other insects, but they may also act as activator repellent in *Aedes* or *Culex* mosquitoes that can detect these odors. It has been proposed that DEET may also ‘confuse’ the olfactory system; this could be tied to its masking effects if its ability to affect volatility varies across odors. While DEET masked all 6 OR ligands we tested, there may be others that are less susceptible to DEET’s effects. This might contribute to olfactory confusion in host-seeking mosquitoes by disrupting sensory input into olfactory circuits underlying mosquito behavioral attraction or host preference^28^.

Our data support the hypothesis that for *An. coluzzii*, synthetic repellents reduce the volatility of OR ligands. This olfactory mode of action may further synergize with effects of these synthetic compounds on other sensory modalities. For instance, recent data in *Aedes aegypti* mosquitoes suggests a non-olfactory based function for DEET as a contact repellent^29^. *Aedes* mosquitoes contain sensory neurons on their tarsi that mediate DEET repulsion. While the DEET-receptor and sensory neurons on the tarsi remain to be identified, they may share a conserved function across many insects. For example, DEET is effective against ticks^30–33^, which do not express Orco or ORs^34^. Interestingly, high concentrations of DEET need to be applied (typically >30%) for it to be effective. Our data suggest this may have two effects. First, we found chemical masking by DEET is most effective at concentrations >30%. Second, as mosquito tarsi are exposed during landing, sufficiently high concentrations of DEET or other insect repellents may be able to trigger contact repellent receptors to elicit repellent behaviors. As such, the effectiveness of DEET against mosquito biting could be due to two overlapping characteristics: its olfactory effect in reducing host-attraction, and its contact effect as a repellent.

Our data suggest that chemicals which reduce the volatility of key host odorants might be effective as host-seeking protectants. An ideal mosquito repellent or repellent mixture might be one that combines three modes of action: active odor-based repellency, odor masking, and contact repellency. Repellents like lemongrass oil were less affected by chemical masking and their combinational use may increase the potency of DEET-based products. Future studies monitoring neural responses directly in the mosquito could yield insights into the function of new repellents as they are identified, as well as streamline the discovery of improved insect repellents.

## Methods

### Mosquitoes

*Anopheles coluzzii* mosquitoes (genotype: *Orco-QF2, QUAS-GCaMP6f*^7^) were raised in a climate chamber maintained at 26-28 °C, 70-80% RH and L14:D10 cycle. After hatching, mosquito larvae were fed on fish food (TetraMin®), added every day. Cotton rolls soaked with sugar solution (10 %, w/vol) were provided to feed adult mosquitoes as a source of carbohydrates. Mosquito females were blood fed on mice for egg laying. The blood feeding protocol was approved by the Johns Hopkins University Animal Care and Use Committee. For calcium imaging, only non blood-fed females (3-10 day old) were used.

### Generation of transgenic *QUAS-GCaMP6f* mosquitoes

#### Cloning of *pXL-BACII-ECFP-15xQUAS-TATA-Gcamp6f-SV40*

The *GCamp6f-SV40-terminator* sequence was PCR amplified from genomic DNA of transgenic *Drosophila* carrying a *QUAS-GCamp6f* transgene (gift from Ya-Hui Chou, unpublished) with primers pBac-TATA-GCamp-SV40-Inf-FOR (5’-gcg gcc gcg gct cga gat ggg ttc tca tca tca tca tc-3’) and pBac-TATA-GCamp-SV40-Inf-REV (5’-ttc aca aag atc gac gtc taa gat aca ttg atg agt ttg gac aaa c-3’). The PCR product was InFusion-cloned (Clontech, catalogue number 639645) into the *pBAC-ECFP-15xQUAS-TATA-SV40* plasmid^7^ (Addgene #104875), digested with ZraI and XhoI. The cloning product was verified by DNA sequencing.

Injections were performed into *Anopheles coluzzii* N’Gousso strain embryos by the Insect Transformation Facility (Rockville, MD) using standard procedures as previously described^7^. Two transgenic lines were established, CP-04-15-M2 and CP-04-15-M3. In functional pilot experiments in crosses to *Orco-QF2* transgenic mosquitoes, both showed similar levels of induced expression and olfactory-directed calcium responses. CP-04-15-M2 was used for all subsequent experiments.

### Mosquito preparation

3-10 day old female mosquitoes were immobilized on ice for 1 min. A mosquito was then carefully inserted into a pipette tip. The mosquito was pushed so only the antennae extended outside the pipette tip. The pipette tip was then attached to a glass slide using modeling clay. For imaging, an antenna was placed forward and flattened on a glass cover slip using two pulled glass capillary tubes (Harvard Apparatus, 1 OD × 0.5 ID × 100 L mm). One tube was used to flatten the 3rd-4th antennal segment, and the other to flatten the 12^th^-13th segment (the most distal segments). Preliminary recordings were performed to visualize responses from the whole antenna. Olfactory responses were similar in each segment but could vary in the number of responding neurons. To achieve higher resolution imaging for analyses, all subsequent recordings were done at one antennal segment (11^th^ antennal segment). Based on pilot experiments examining multiple segments, the responses in one segment (11^th^ segment) were representative of responses in all segments.

### Odorants

DEET, *p*-cresol, and 2-acetylthiophene were purchased from Sigma-Aldrich. Eugenol, benzaldehyde, and Indole were purchased from Aldrich. Lemongrass oil, 1-octen-3-ol, and 1-hepten-3-ol were purchased from SAFC. IR3535 was purchased from EMD Chemicals. Picaridin was purchased from Cayman Chemical. Repellents were either used undiluted, or diluted to 1%, 10%, or 30% in paraffin oil (Sigma-Aldrich). OR ligands were diluted to 1% in paraffin oil.

### Odorant delivery

For testing neural responses towards OR ligands and repellents, 20 μl of the solution was pipetted onto a piece of filter paper (1×2 cm) placed in a Pasteur pipette. For mixtures, 10 μl of an OR ligand was pipetted along with 10 μl of repellent on the same filter paper. Each odorant was prepared at double the final concentration to reach the desired final concentration when mixed. The Pasteur pipette was then inserted into a hole in a plastic pipette (Denville Scientific Inc, 10ml pipette) that carried a purified continuous air stream (8.3 ml/s) directed at the antenna. A stimulus controller (Syntech) was used to divert a 1 s pulse of charcoal-filtered air (5 ml/s) into the Pasteur pipette starting 10 seconds after the beginning of each recording. Each animal was tested with 6 odorant pairs (6 OR ligands and their respective mixtures). Four animals out of a total of 45 animals stopped responding before testing all odorants, and the remaining odorant pairs were tested in new animals. The sequence of odorants was randomized, and recordings from a mosquito were discarded if a response to a positive control odorant (usually 1-octen-3-ol) was absent. New Pasteur pipettes were prepared for each recording day.

### Modified odorant delivery

To test whether masking occurs at the receptor or the chemical level, the odorant delivery described above was modified by three different methods:

1. Pre-stimulation with repellents: An OR ligand (1-octen-3-ol) and a repellent (DEET, IR3535, picaridin, or paraffin oil for control) were prepared in two separate Pasteur pipettes as previously described. Each Pasteur pipette contained 10 μl of either 2% 1-octen-3-ol or 60% repellent to reach a final concentration of 1 and 30%, respectively. The two Pasteur pipettes were inserted into two holes in the plastic pipette that carried a purified continuous air stream directed at the antenna. One branch of a polyethylene Y-tube was used to deliver a 3 s pulse of charcoal-filtered air into the Pasteur pipette that contains the repellent. At the third second, the other branch of the Y-tube was attached to the 1-octen-3-ol Pasteur pipette to deliver 1 s pulse of1-octen-3-ol. For comparison, a mixture of the repellent and 1-octen-3-ol was also tested with each animal as previously described. Each animal was tested with 7 odorant conditions.
2. Simultaneous odorant delivery: an OR ligand (1-octen-3-ol) and a repellent (or paraffin oil for control) were prepared in two separate Pasteur pipettes as previously described. The two Pasteur pipettes were inserted into two holes in the plastic pipette that carried a purified continuous air stream directed at the antenna. A 1 s pulse of charcoal-filtered air (5 ml/s) was diverted into the two Pasteur pipettes using a polyethylene Y-tube in order to deliver the two odorants at the same time into the continuous air stream. Afterwards, the two Pasteur pipettes were switched between the two holes in the long plastic pipette to rule out any position bias. For comparison, a mixture of the repellent and 1-octen-3-ol was also tested with each animal as previously described. Each animal was tested with 11 odorant conditions.
3. Same pipette delivery: an OR ligand (1-octen-3-ol) and a repellent (or paraffin oil for control) were applied on two separate filter papers (0.5×1 cm) within the same Pasteur pipette. We made certain the two filter papers were not touching and therefore the odorants were never physically mixed. To deliver the odorants, a 1 s pulse of charcoal-filtered air was diverted into the Pasteur pipette. Afterwards, we used another Pasteur pipette, in which the position of the repellent and 1-octen-3-ol was swapped, to rule out any position bias. For comparison, a mixture of the repellent and 1-octen-3-ol was also tested with each animal as previously described. Each animal was tested with 11 odorant conditions.

### Imaging system

Antennae were imaged through a 10x (Zeiss EC Epiplan-Neofluar 10x/0.25) and a 50x (LD EC Epiplan-Neofluar 50x/0.55 DIC) objectives mounted on a Zeiss Axio Examiner D1 microscope. For fluorescence, a light source (Zeiss Illuminator HXP 200C) and eGFP filter cube (FL Filter Set 38 HE GFP shift free) were used.

For image acquisition, an EMCCD camera (Andor iXon Ultra) and NIS Elements Advanced Research software (Nikon instruments) were used. Recordings were for 20 seconds, at a resolution of 512×512 pixels, and an exposure time of 200 ms (5 Hz).

### Analysis of Calcium imaging recordings

To make the heatmap ΔF images, a custom-built macro in Fiji was used. This Macro uses the “Image stabilizer” plug-in to correct for movements in the recording, followed by the “Z project” function to calculate the mean baseline fluorescence (mean intensity in the first 9 seconds of recording, before stimulus delivery). Then, the “Image calculator” function was used to subtract the mean baseline fluorescence from the image of maximum fluorescence after odorant delivery (this image was manually chosen). Afterwards, this ΔF image was used to produce heatmaps.

To produce intensity time traces, the “ROI manager” tool in Fiji was used to manually select ROIs. ROIs were drawn around cells that showed increased fluorescence (based on the ΔF image). Then the “multi-measure” function in the “ROI manager” was used to produce intensity values for those ROIs across time. Finally, these values were saved into Excel and used to calculate *ΔF/F*\*100. *ΔF/F*\*100 = *F*_i_/*F*_0_*100, where *F*_i_ is the fluorescence intensity value at frame _i_, while *F*_0_ is the mean fluorescence intensity before odorant delivery (first 9 seconds, 45 frames).

Maximum responses (Maximum *ΔF/F*\*100 values) were used in the following analysis.

Linear Mixed Effects (LME) regression was used to model the average value of the outcome under an experimental condition, accounting for both correlation due to repeated measurements and non-constant residual variation. In all experiments, fixed effects were used to model the average value of the outcome at each experimental condition, and a linear term was used to model the average change in the outcome over repeated measurements. Within-subject correlation was accounted for using random intercepts, and heteroskedasticity was accounted for by modeling the residual variance.

For odorant delivery and pre-stimulation experiments, the residual variance was modeled as a power of the fitted values. In the simultaneous odorant delivery experiments, the outcome was log transformed and a separate residual variance term was estimated for conditions where repellents were physically mixed with the OR ligands. In the same pipette delivery experiment, the outcome was log transformed and the residual variance was modeled as an exponential function of fitted values.

Model assumptions, such as linearity of relationships, normally distributed scaled residuals, and normally distributed random effects, were assessed using residual diagnostic plots. Confidence intervals and p-values provided use the Wald approximation. No multiple comparisons corrections were performed. All analyses were performed using R version 3.5.1^35^ using the nlme package version 3.1-137^36^.

### Photoionization detector

The MiniRAE 3000 photoionization detector (Honeywell RAE Systems) was used to calculate concentrations of odorants delivered to the mosquito antenna in different experiments. The photoionization detector was calibrated to a reference gas (ethyl acetate) and was attached to the tip of the plastic pipette used to deliver odorants in calcium imaging experiments. The maximum reading (concentration in ppm) following each odorant delivery was reported.

## Acknowledgements

We thank C. McMeniman, D. Task, S. Maguire, and K. Robinson for mosquito technical support and for comments on the manuscript; Sophia Hager for assistance with imaging experiments; Mark Wu for the use of his imaging camera. This work was supported by grants from the Department of Defense to C.J.P. (W81XWH-17-PRMRP), from the National Institutes of Health to C.J.P. (NIAID R01Al137078), a Johns Hopkins 2018 Catalyst Award to C.J.P., a Johns Hopkins Malaria Research Institute Pilot Fund to C.J.P., and a Johns Hopkins Malaria Research Institute Postdoctoral Fellowship to A.A.

## Author Contributions

A.A. and C.J.P. conceived the study and experimental design. A.A. carried out all experiments. A.A. analyzed the calcium imaging data. J.B. performed statistical analyses on the calcium imaging data. O.R. generated the *Anopheles* strain containing *QUAS-GCamP6f*. A.A. and C.J.P wrote the paper with input from O.R.

## Data Availability

The imaging files and datasets generated during and/or analysed during the current study are available from the corresponding author on reasonable request.

---

**Figure S1.**
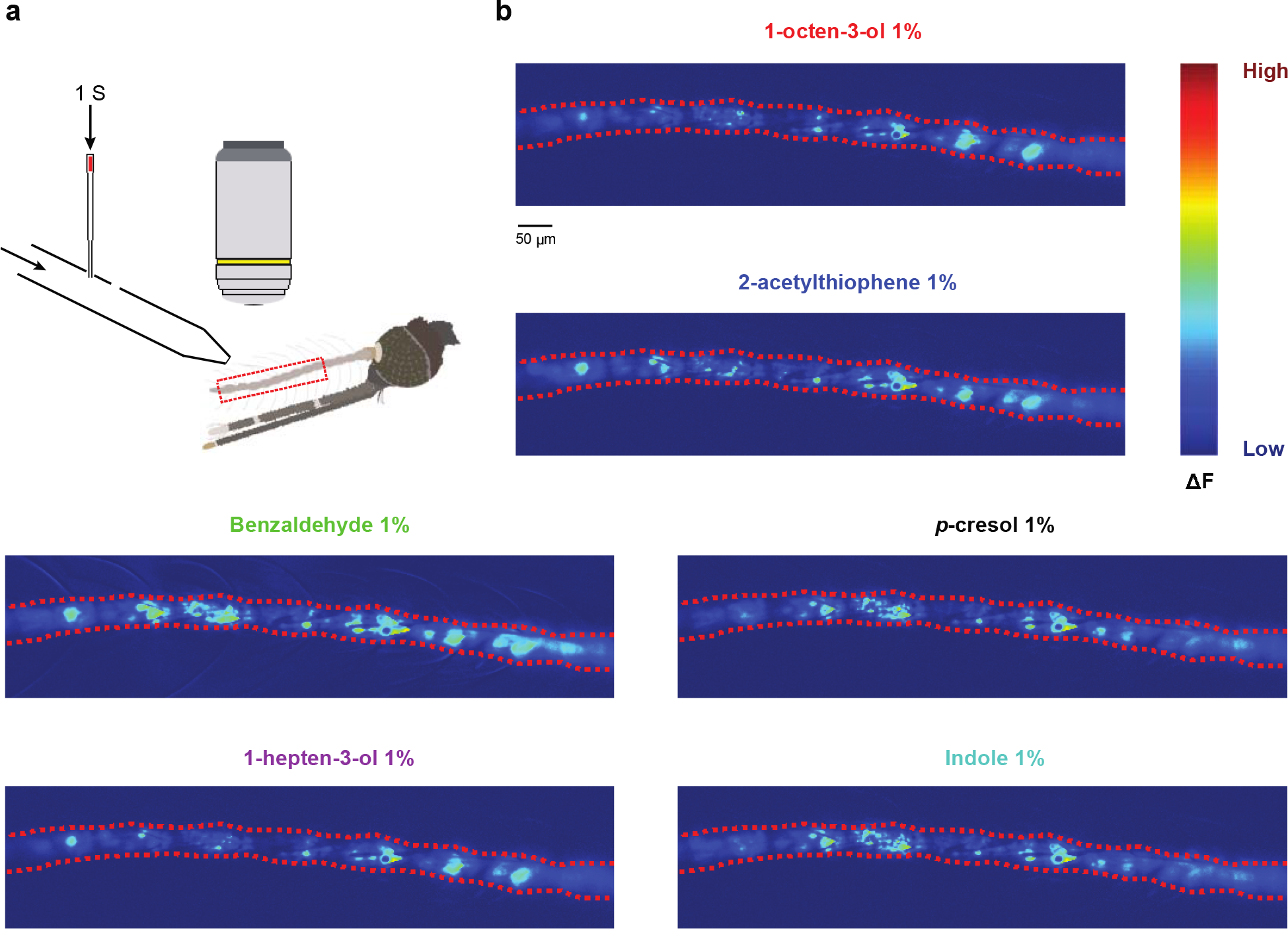
OR ligands globally activate *An. coluzzii* antennal Orco expressing olfactory neurons. **a,** Schematic of the calcium imaging setup. A 10x microscope objective is used to image most of the antenna at a time (∼ 9/12 olfactory antennal segments outlined in dashed red rectangles). Arrows indicate the direction of air flow (continuous air, and 1 s air pulse). **b,** Heatmaps of the responses towards each OR ligand at 1% (dashed red lines indicate the borders of the antennae). ΔF values are arbitrary units.

**Figure S2.**
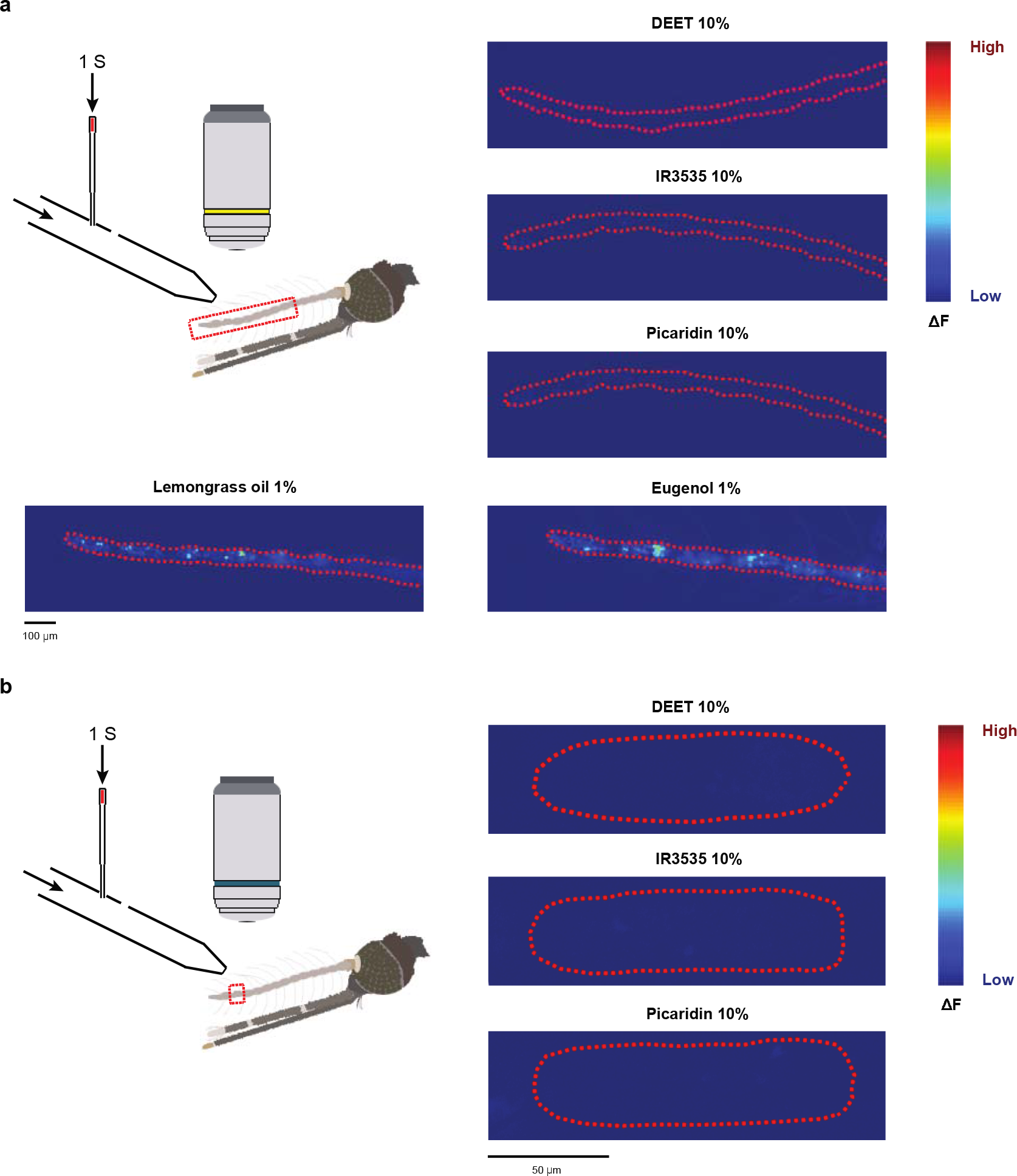
Natural repellents, but not synthetic repellents, strongly activate *Anopheles* olfactory neurons. **a,** Responses across most of the antenna (∼ 9 segments, dashed red line) towards 10% DEET, 10% IR3535, and 10% picaridin, and towards 1% of lemongrass oil and 1% eugenol. **b,** Higher magnification responses at the 11^th^ antennal segment (dashed red line) towards 10% DEET, 10% IR3535, and 10% picaridin.

**Figure S3.**
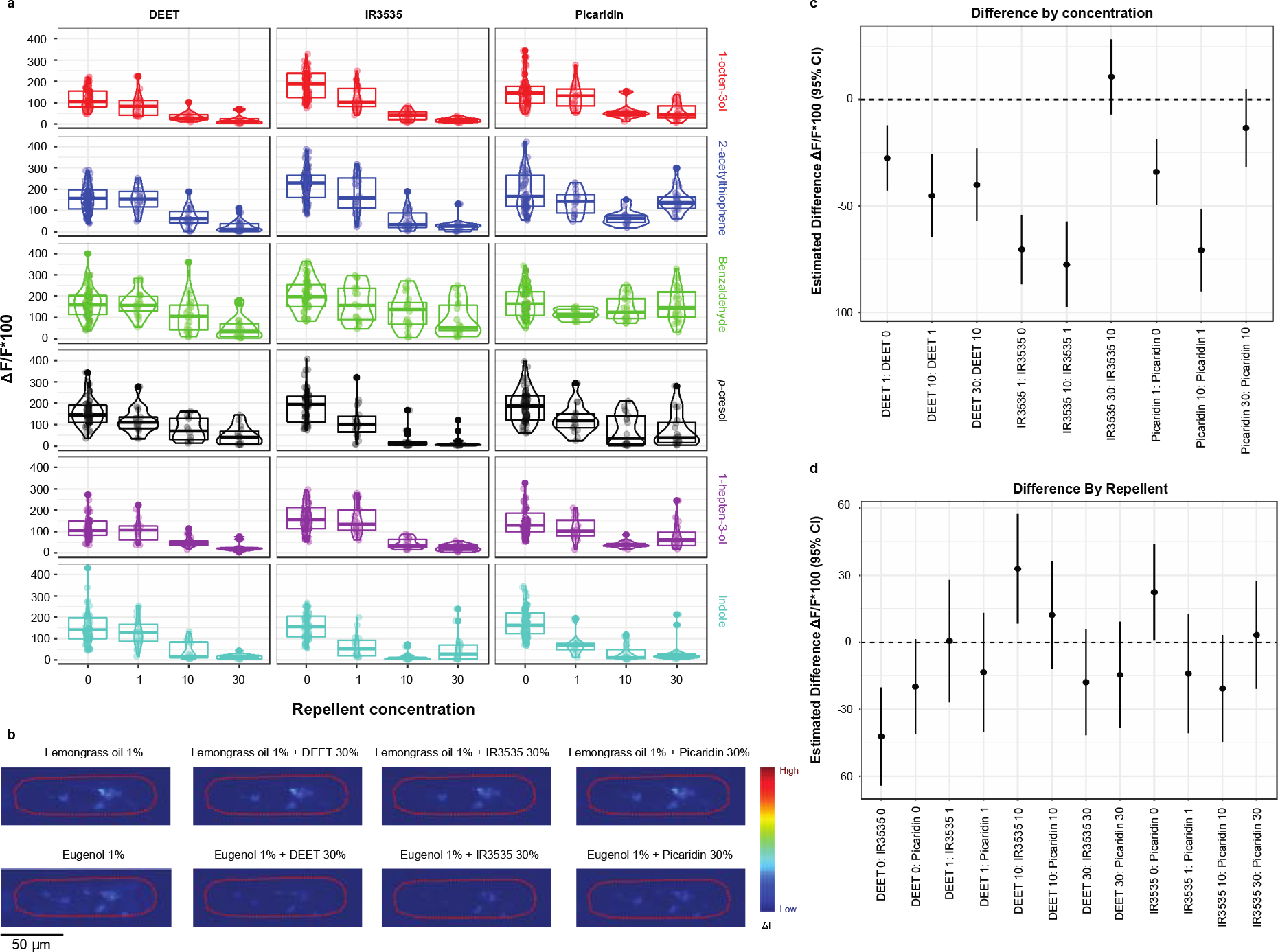
DEET, IR3535, and picaridin mask the olfactory responses towards odorants. **a,** Box plots (median and 25^th^-75^th^ percentiles) representing raw data for the olfactory responses towards mixtures of the six OR ligands at 1% with repellents (DEET, IR3535, and picaridin) at 0% (OR ligand alone), 1%, 10%, and 30% concentrations (n=15 animals for each condition of 0% repellent, n=5 animals for all other conditions, 1-7 responding neurons/animal, each dot represents a responding neuron). **b,** Heatmaps of the responses towards 1% of activator repellents (lemongrass oil and eugenol) and their mixtures with 30% DEET, 30% IR3535, or 30% picaridin (n=5 animals for each condition, 2-4 responding neurons/animal). **c,** LME contrasts estimates (mean and 95% Wald CI) between subsequent concentrations of the same repellent. **d,** LME contrasts estimates (mean and 95% Wald CI) between different repellents at the same concentration. For c and d, CI intersecting with the dashed 0 line indicate non-significant difference (P > 0.05, LME model with Wald approximation).

**Figure S4.**
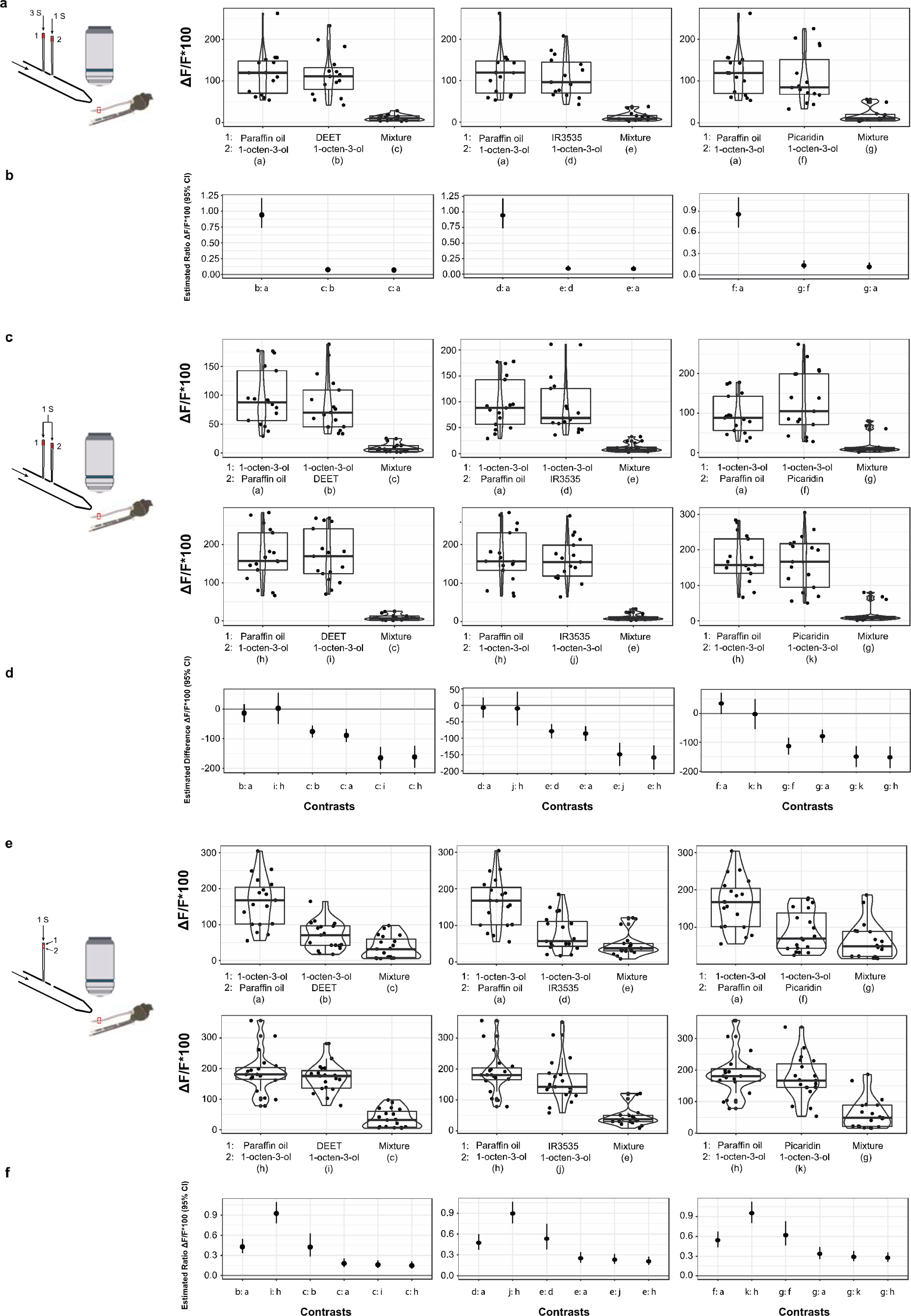
Olfactory responses to OR ligands are decreased when mixed with synthetic repellents. **a,** Box plots (median and 25^th^-75^th^ percentiles) representing raw data for the responses towards a 1 s pulse of 1% 1-octen-3-ol occurring during the last second of a 3 s pulse of paraffin oil, 30% DEET, 30% IR3535, or 30% picaridin, compared to the response towards physical mixtures of 1% 1-octen-3-ol with 30% of DEET, 30% IR3535, or 30% picaridin. The numbers next to odorant names indicate the position of the Pasteur pipette that contains that odorant in the schematic. **b,** LME contrasts estimates (mean and 95% Wald CI) between odorant conditions in a (letters on the X-axis correspond to the letters in a to indicate odorant conditions). **c,** Box plots (median and 25^th^-75^th^ percentiles) representing raw data for the responses towards a 1 s pulse of 1% 1-octen-3-ol in the first position or the second position simultaneously delivered with a 3 s pulse of paraffin oil, 30% DEET, 30% IR3535, or 30% picaridin, compared to the response towards physical mixtures of 1% 1-octen-3-ol with 30% of DEET, 30% IR3535, or 30% picaridin. **d,** LME contrasts estimates (mean and 95% Wald CI) between odorant conditions in c (letters on the X-axis correspond to letters in c to indicate odorant conditions). **e,** Box plots (median and 25^th^-75^th^ percentiles) representing raw data for the responses towards a 1 s pulse of 1% 1-octen-3-ol when applied on the upper filter paper or the lower filter paper with paraffin oil, 30% DEET, 30% IR3535, or 30% picaridin in the same Pasteur pipette, compared to the response towards physical mixtures of 1% 1-octen-3-ol with 30% of DEET, 30% IR3535, or 30% picaridin. **f,** LME contrasts estimates (mean and 95% Wald CI) between odorant conditions in e (letters on the X-axis correspond to letters in e to indicate odorant conditions). Contrasts in b and f come from a log transformed model, representing a ratio of geometric means: *e.g.* a value of 1 indicates no difference, a value of 0.5 indicates a 50% reduction, and a value of 1.5 indicates a 50% increase. Contrasts in d come from a model fit to the raw data; CI intersecting with the dashed 0 line indicate non-significant difference. For a,c,e, n=5 animals for each condition, 1-6 responding neurons/animal, each dot represents a responding neuron.

